# Intrinsically Disordered Regions Define Unique Protein Interaction Networks in CHD Family Remodelers

**DOI:** 10.1101/2024.08.25.609601

**Authors:** Mehdi Sharifi Tabar, Chirag Parsania, Caroline Giardina, Yue Feng, Alex CH Wong, Cynthia Metierre, Rajini Nagarajah, Bijay P Dhungel, John EJ Rasko, Charles G Bailey

**Author notes:** Contributed equally (Chirag Parsania, Caroline Giardina).

## Abstract

Chromodomain helicase DNA-binding (CHD1-9) enzymes reposition nucleosomal DNA for transcription, recombination, and replication. They possess highly conserved ATPase domains flanked by poorly characterised N- and C-termini, which are enriched with intrinsically disordered regions (IDRs) and short aggregation-prone regions (APRs). The roles of IDRs and APRs in CHD function has remained elusive. Here, by integrating proteomics and AlphaFold Multimer analysis, we defined the protein-protein interaction (PPI) networks within the N- and C-termini of all CHDs. We generated a comprehensive map of CHD1-9-specific binding proteins, revealing dozens of novel interactions with transcription regulators. We identified APR regions that contribute to PPI formation and demonstrated that a highly conserved APR within the C-terminus of CHD4 is critical for its interaction with the nucleosome remodeling and deacetylase (NuRD), as well as the CHD, ADNP, and HP1 (ChAHP) complexes. Further analysis unravels a regulatory role for the CHD4 APR in gene transcription during erythrocyte formation. Our results emphasize that the N- and C-termini of CHD chromatin remodelers establish PPI networks that drive unique transcriptional programs.

## Introduction

In eukaryotes, the basic unit of chromatin is the nucleosome, which consists of 147 bp of DNA wrapped around an octamer of histone proteins(Bednar *et al*, 1998). The nucleosomal structure restricts the DNA accessibility of various DNA processing complexes involved in transcription, replication, and repair(Lai & Pugh, 2017; Saha *et al*, 2006). The DNA helicase/ATPase domain of chromatin remodeler proteins hydrolyses ATP to distort histone-DNA interactions and thereby control the accessibility of DNA to various DNA processing complexes(Clapier *et al*, 2017; Farnung *et al*, 2020; Hargreaves & Crabtree, 2011; Zhong *et al*, 2020).

Chromatin remodelers are categorised into different families that include SWI/SNF, ISWI, INO80, and CHD protein members(Clapier *et al*., 2017). The CHD family remodelling proteins are mainly involved in nucleosome assembly and organisation during replication and transcription. They are categorised into three subfamilies based on their sequence identity, domain architecture and functional properties: subfamily I (CHD1-2), subfamily II (CHD3-5); which are found in the NuRD and the ChAHP complexes(Low *et al*, 2020; Ostapcuk *et al*, 2018; Sharifi Tabar *et al*, 2022a; Sharifi Tabar *et al*, 2019); and subfamily III (CHD6– 9)(Marfella & Imbalzano, 2007; Trujillo *et al*, 2022). Only CHD1 is found in yeast and the expansion of the CHD family in higher eukaryotes allows for more specialised and finely tuned regulation of chromatin dynamics and gene expression. The interaction of remodelling proteins with various chromatin binding factors expands their capacity to fine-tune gene expression programs in many biological processes. However, owing to the large size of CHD proteins and their high affinity for nucleosomal DNA, protein interaction studies have been technically challenging. The rigorous identification of new binding partners will facilitate characterisation of biological functions within and between CHD subfamilies which may occur in a tissue- or context-specific manner.

All CHD proteins belong to the superfamily 2 (SF2) ATPase family, which consists of two lobes: DExx and HELICc(Durr *et al*, 2006; Enemark & Joshua-Tor, 2008). These lobes form an active-site cleft for ATP hydrolysis that produces the required energy for DNA translocation. Additionally, all CHD members contain tandemly arranged chromatin organisation modifier (chromo) domains located before the ATPase domain, which are primarily involved in binding methylated lysine residues on histones. The central chromo and helicase domains of CHDs are flanked by poorly characterised N- and C-termini that contain aggregation prone regions (APRs) and/or intrinsically disordered regions (IDRs). These regions engage in both transient and stable protein interactions and mediate phase separation of chromatin remodellers in various biological processes(Patil *et al*, 2023). Here, we utilised affinity purification followed by mass spectrometry (AP-MS) to map the mutual and unique interaction partners of the N- and C-termini of all CHD members(Sharifi Tabar *et al*, 2022b). We identified dozens of novel interactions that are either unique or common to different CHD proteins. Using AlphaFold Multimer we assessed the structural basis of several candidate interactions. By integrating our data from AP-MS and AlphaFold Multimer, we identified and characterized a CHD4-specific APR that can mediate interaction of CHD4 with the nucleosome remodelling and deacetylase (NuRD) and CHD, ADNP, HP1 (ChAHP) complexes in a mutually exclusive manner.

## Results

### CHD N- and C-termini are highly disordered

Both the chromo and ATPase domains of CHD proteins are centrally located and flanked by mostly unstructured N- & C-termini containing unique functional domains (**Fig 1.A**). Only subfamily II (CHD3-5) N-termini contain a high mobility group (HMG) Box-like domain and tandem plant homeodomains (PHDs) that bind to poly(ADP-ribose) and H3K9me3 post-translational marks, respectively(Silva *et al*, 2016) (**Fig 1.A**), whereas the C-termini contain a poorly defined CHD C-terminal (CHDCT2) domain. Other C-terminal domains include a structured domain of unknown function (DUF) in CHD1 and Brahma and Kismet (BRK) domains in members of CHD subfamily III (**Fig 1.A**).

**Figure 1.**
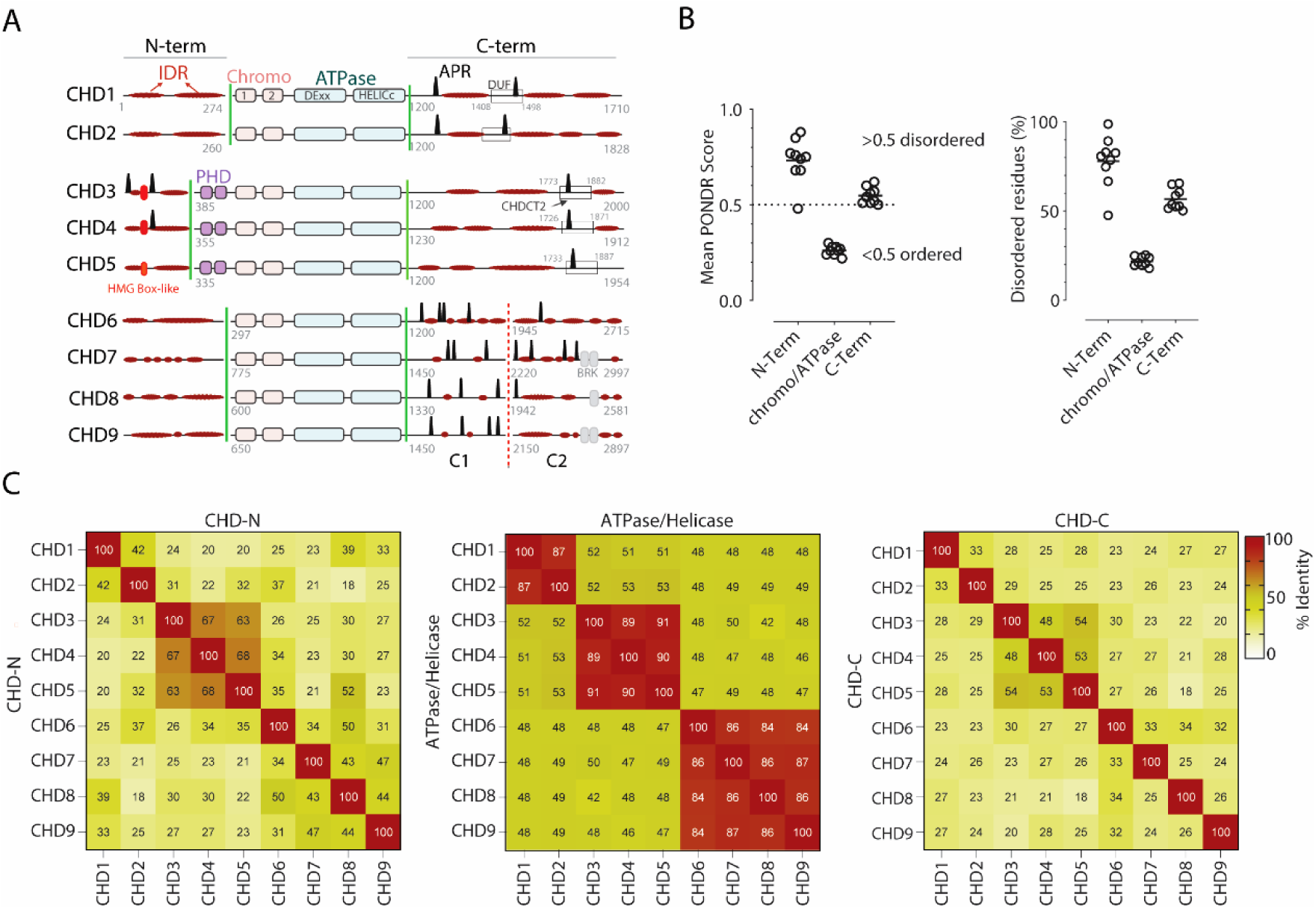
CHD family members exhibit both unique and shared structural characteristics. **A)** Representation of the domain organization of CHD proteins, highlighting the distribution of aggregation prone regions (APRs, black peaks) and intrinsically disordered regions (IDRs, flat red ovals) within the N- and C-termini. The domain of unknown function (DUF) and CHD C-terminal 2 (CHDCT2) domains, which contain APR regions within the C-termini, are represented by rectangles. Green lines denote boundaries for N- and C-terminal segments used in AP-MS analysis. The C1 and C2 segments, separated by a dotted red line, denote further truncated CHD6-9 C-termini. Numbers in grey refer to amino acid positions. Red and grey ovals represent high mobility group (HMG) Box-like and brahma and kismet (BRK) domains, respectively. **B)** The graphs illustrate the mean PONDR score and percentage disordered residues of the N-terminal, ATPase, and C-terminal domains of all CHD proteins. **C)** Heatmap matrices represent Pearson correlation analysis within and between CHD subfamily domains.

We examined the presence of IDRs within the N- and C-termini which revealed long stretches of highly disordered residues (**Fig 1.B**). We used PONDR, a web-based computational tool, to predict the probability of disordered regions in all CHDs. PONDR scores confirmed that both the N- and C-termini are highly disordered (score >0.5), while the ATPase domains are highly ordered (score <0.5) (**Fig 1.B**). Additionally, about 80% of the N-terminal residues and 53% of the C-terminal residues are disordered, whereas only 21% of the ATPase domain residues are disordered (**Fig 1.B**). We also examined the number and distribution patterns of APRs in CHD proteins using TANGO, a web-based computer algorithm for prediction of aggregating regions in unfolded polypeptide chains (**Fig 1.A**). While the N-termini of subfamilies I and III are devoid of APRs, only CHD3 and CHD4 members of subfamily II contain 2 and 1 APRs within the N-termini, respectively (**Supplementary Table 1**). C-terminal analysis showed that subfamilies I and III contain 2 or more APRs, whereas only subfamily II members possess a unique APR within their CHDCT2 domain (**Fig 1.A**). In the central ATPase domain-containing region of CHDs our sequence alignment analysis reveals that there is ∼87% sequence identity across various CHD proteins within each subfamily (**Fig 1.C**). In contrast, sequence alignment and identity analysis of the N- and C-termini of CHD family proteins indicates that they are much less conserved with sequence identity ranging from 24-54% (**Fig 1.C**).

To map the CHD termini-specific protein interaction networks, we cloned the N- and C-termini of all CHDs into a mammalian expression plasmid also containing a FLAG epitope (**Fig 1.A**). The longer C-termini of CHD6-9 were divided into two distinct segments, identified as C1 and C2, to facilitate expression. (**Fig 1.A**). All fragments were transiently overexpressed in Expi293 cells, the nuclear fraction was collected, and the FLAG-tagged CHD fragments were immunoprecipitated using FLAG antibody-conjugated beads. To ensure that all fragments were expressed and purified we evaluated the abundance of FLAG-CHD by MS. Of 22 CHD fragments tested, 19 were detected and quantified (**Supplementary Fig 1**). Fragments that did not show any detectable expression (CHD7-N, CHD6-C1 & 2) or failed to immunoprecipitate any proteins (CHD8-C1) were excluded from further analysis.

### The protein interactions of N-termini

AP-MS analysis of CHD N-termini fragments revealed an enrichment of chromatin binding or modifying factors (**Fig 2, Supplementary Table 2**). A distinct protein interaction signature within and between some CHD proteins and subfamilies was observed (**Fig 2**). For instance, the general TAT-binding associated factors (TAF1B, TAF1C, TAF1D) were enriched with CHD1. TAFs nucleate RNAPII formation at promoter regions and CHD1 regulates RNAPII initiation and transcription(Joo *et al*, 2017; Lin *et al*, 2011). The interaction might provide functional synergy to control productive transcription. The euchromatic histone lysine methyltransferase (EHMT1/EHMT2) complex, responsible for adding methyl groups to H3K9 histone residues(Brejc *et al*, 2017; Dillon *et al*, 2005), was enriched with the N-termini of various CHDs, including CHD1-2 and CHD6. A marked enrichment of round spermatid basic protein (RSBN1) with CHD1-3 and CHD8 was observed. RSBN1, also known as KDM9, acts as a demethylase on the histone mark H4K20me2, a DNA replication mark(Brejc *et al*., 2017). This provides supportive evidence for the involvement of the remodelers in coordinated gene transcription and DNA replication. We also observed the enrichment of the casein kinase II complex subunits(Borgo *et al*, 2021)(CSK2B, CSK21, and CSK22) with the N-termini of CHD3-4. Other notable enrichments included bromodomain-containing protein (BRD8) with CHD5, acid nuclear phosphoprotein (ANP32A/B) with CHD6, and zinc finger MYM-type-containing (ZMYM2) with CHD9, each of which is known to play roles in chromatin organization and gene expression.

**Figure 2.**
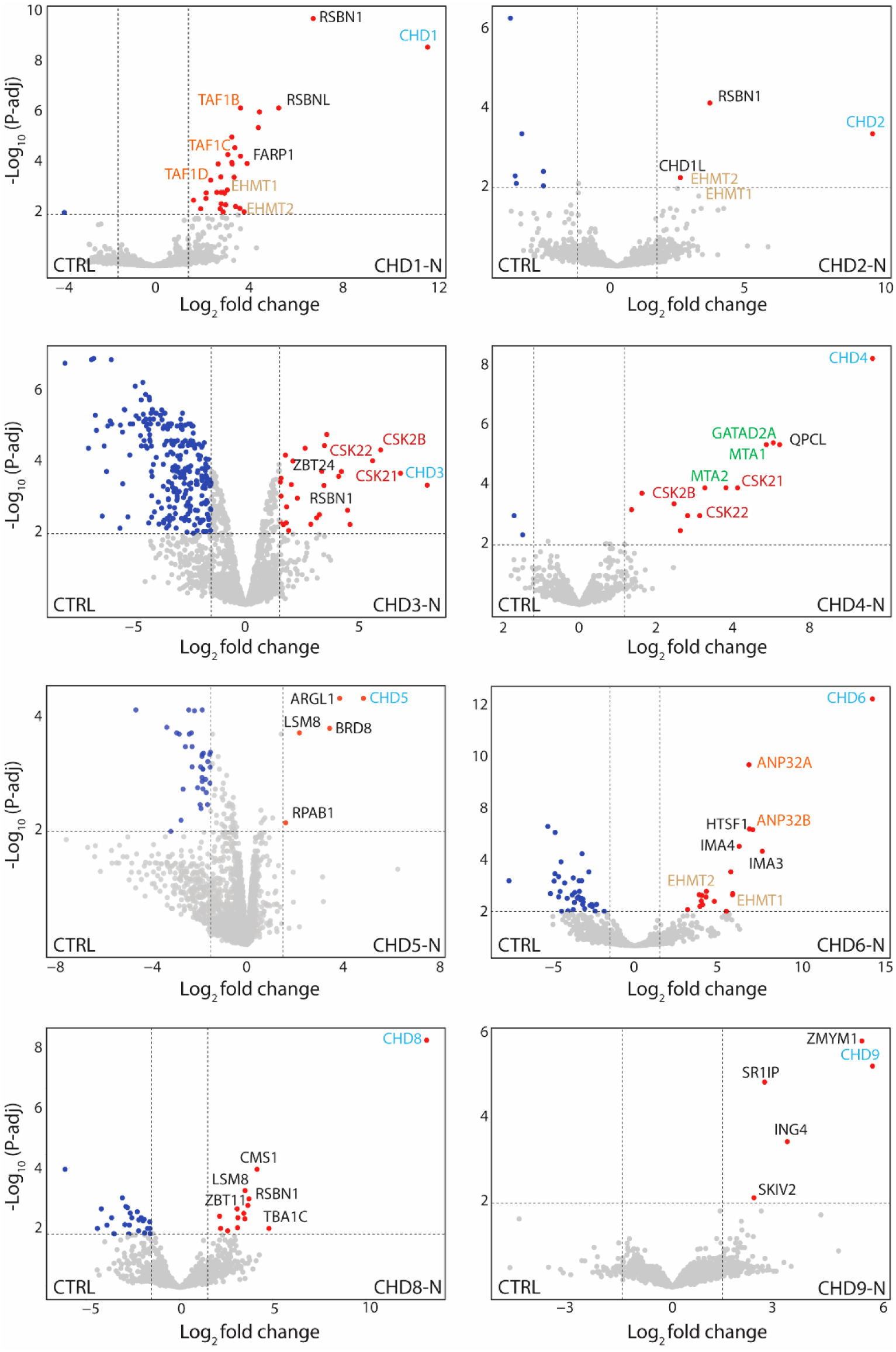
The protein interaction landscapes of CHD N-termini. Volcano plots representing enriched proteins by FLAG pulldowns (n=3). CHD bait proteins are labeled in blue, while co-purified proteins are in black. Subunits of complexes enriched with CHD proteins are color-coded with the same color. Significance was determined as Log_2_(Fold Change) >1.5, - log_10_ (p-adj)> 2; grey dots indicate proteins which were not significantly enriched.

### The protein interactions of C-termini

Analysis of CHD protein C-terminal fragments identified additional unique and shared proteins involved in gene regulation and chromatin remodelling (**Fig 3, Supplementary Table 2**). For example, the enrichment of barrier to autointegration factor (BAF), a core component of the SWI/SNF remodelling complex, with CHD1 is intriguing as both have been shown to localise to the promoter and enhancers of target genes(Brahma & Henikoff, 2024; Jeronimo *et al*, 2021). In addition, histone chaperones including structure-specific recognition protein (SSRP1) and SPT16 homolog (SP16H), subunits of the FACT (facilitates chromatin transcription) complex, were enriched with the CHD1 C-terminus. Together, CHD1 and FACT facilitate Pol II passage through nucleosomes to aid transcription(Farnung *et al*, 2021). Our observations reveal the co-enrichment of RTF1, an RNAPII-associated protein, further validating previous findings and underscoring its pivotal role in transcriptional regulation(Simic *et al*, 2003) (**Fig 3**). Consistent with our previous findings, we observed a marked enrichment of the NuRD complex subunits and activity-dependent neuroprotective protein (ADNP), the core subunit of the ChAHP complex, with the C-termini of CHD3-5(Sharifi Tabar *et al*., 2022a; Sharifi Tabar *et al*., 2019; Torrado *et al*, 2017). A notable enrichment of KIAA1671, exclusively with the C-termini of CHD3-5, but no other CHDs (**Fig 3**), may suggest a unique role for this poorly characterised protein in gene regulation and chromatin organisation. ZMYM1, a lowly expressed nuclear protein, was specifically enriched with the C-terminus of CHD9. DNAJ homolog subfamily B member, DNAJB6, was co-enriched with the C-termini of most CHD proteins, which suggests a possible transcriptional regulatory role for this chaperone as a stimulatory effector on ATPase activity. In summary, our analysis provides an important resource comprising both unique and shared proteins that bind via the N- and C-termini of CHD family proteins.

**Figure 3.**
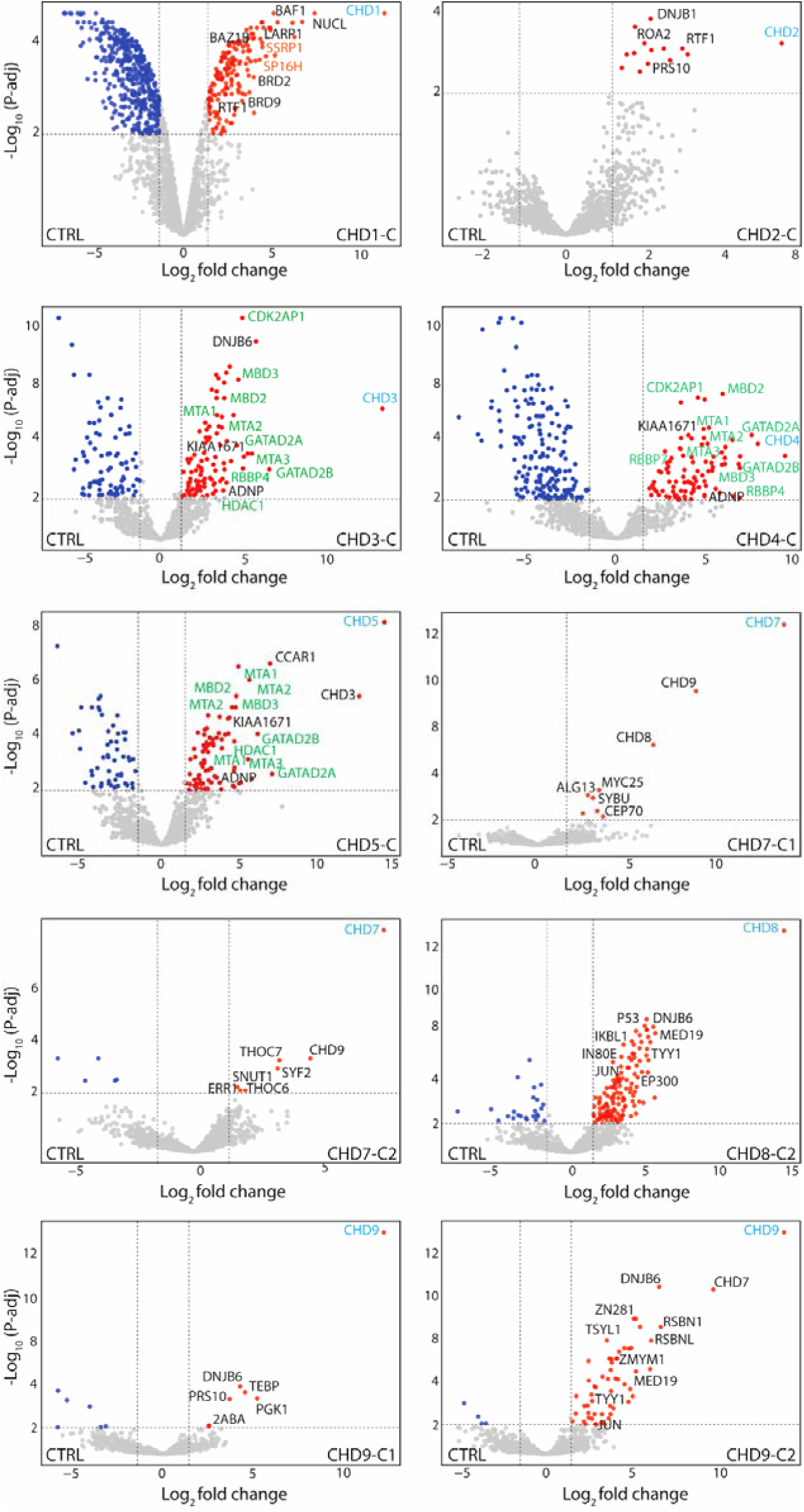
The protein interaction landscapes of CHD C-termini. Volcano plots representing enriched proteins by FLAG pulldowns (n=3). CHD bait proteins are labelled in blue and co-purified proteins are in black. Subunits of complexes enriched with CHD proteins are color coded. Significance was determined as Log2(Fold Change) >1.5, -log10 (p-adj)> 2; grey dots indicate proteins which were not significantly enriched.

### Validation of CHD protein interactions with AlphaFold

To independently evaluate the interaction likelihood and structural basis of identified protein interactions, we utilised Alphafold Multimer, an extension of AlphaFold2(Evans *et al*, 2022). We first tested several potential proteins that exhibited strong co-enrichment patterns with CHD proteins. These include the CHD1-N:EHMT1, CHD5-N:BRD8, CHD6-N:ANP32A, CHD9-C2:DNBJ6, CHD2-C:RTF1, CHD4-C:ADNP, and CHD4-C:GATAD2B complexes. High confidence interactions (markedly reduced predicted alignment error (PAE) scores (see Methods)) were obtained between potential interacting domains in the CHD1-N:EHMT1, CHD6-N:ANP32A complexes (**Supplementary Fig 2. A, B**). It is notable that in nearly all these heterodimer complexes, poor pLDDT scores (see Methods) were predicted for CHD polypeptides as these are highly disordered. Low confidence interactions (mildly reduced PAE scores) were obtained for potential interacting domains in the CHD5-N:BRD8 and the CHD9-C2:DNJB6 complexes(Abramson *et al*, 2024; Ruff & Pappu, 2021).

AlphaFold Multimer analysis predicted high confidence (low PAE) interactions between high pLDDT domains for potential heterodimers CHD2-C:RTF1, CHD4-C:GATAD2B, and CHD4-C:ADNP complexes (**Fig 4.A & B**). This is mainly because CHD2-C and CHD4-C contain relatively well structured but functionally poorly understood DUF, and CHDCT2 domains, respectively. Investigating the interaction interfaces for all these three complexes revealed the presence of several α-helical structures including APRs (shown by dark blue circles) at the interaction interfaces (**Fig 4.B**). AlphaFold predicted two potential interaction interfaces for the CHD2-C-RTF1 complex. In this complex, the CHD2-C-APR region (residues 1477-1488) can potentially interact with residues 61-79 and/or 301-311 on the RTF1 protein. For the CHD4-C-GATAD2B complex, AlphaFold predicted two potential regions on CHD4-C: the first region includes residues 1453-1513, and the second region includes residues 1731-1744, which encompass the APR region. These regions might interact with either residues 372-404 on the GATAD2B protein or residues 9-41 on the ADNP protein.

**Figure 4.**
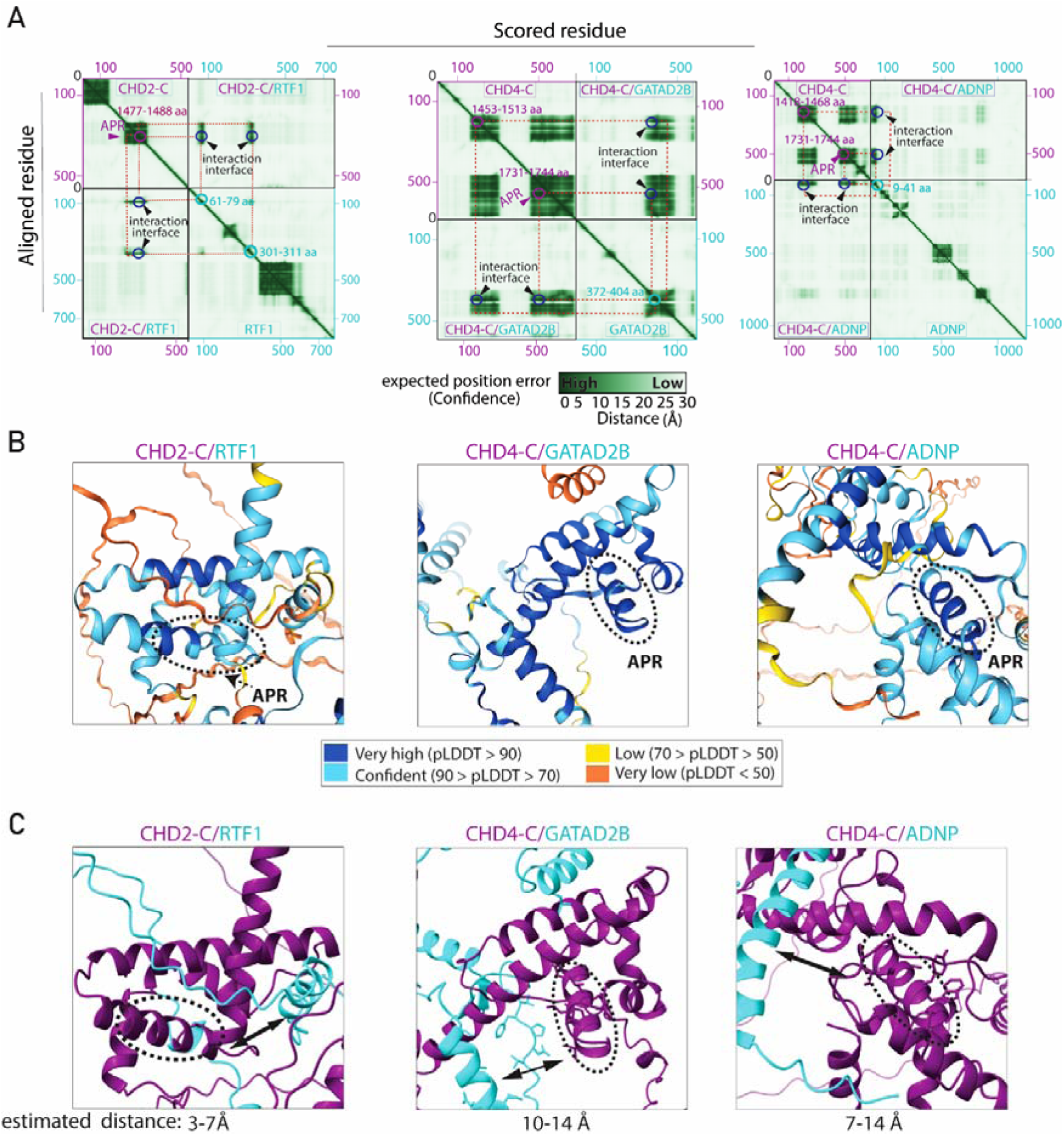
AlphaFold validation of CHD protein interactions discovered by AP-MS. **A)** Heatmaps represent the Predicted Aligned Error (PAE) score between all pairs of residues. A dark green color indicates a low PAE, signifying high reliability of the relative position of the residues. Conversely, lighter shades denote lower confidence in the positioning. Potential interaction interfaces are indicated by arrowheads. Potential interacting residues within each polypeptide are indicated in the same color as the protein label. **B)** Predicted 3D structure of the identified complexes. The predicted local distance difference tool (pLDDT) score represents the confidence of the predicted structure. APRs form alpha helical structures with high pLDDT scores (highlighted with dotted circles in B & C). **C)** Estimated contact distances between APRs (purple) and interacting proteins (cyan); left, between CHD2-C and RTF1; middle, between CHD4-C and GATAD2B; right, between CHD4-C and ADNP. Double-headed arrows in C indicate the estimated distance between the two polypeptides.

To investigate whether APRs are involved in the direct interaction with RTF1, GATAD2B, and ADNP, we measured the distance between APRs and the closest residues of the interacting partner using Chimera X. Our analysis suggests that the CHD2-C interaction with RTF1 could be direct, with an estimated distance of 3-7Å. Measurement of CHD4-C-specific APRs distance with both ADNP and GATAD2B suggest a distance greater than 7 Å. This highlights that CHD4-C APR may not be directly involved in the interaction, however, it does not exclude the contribution of the APR in protein interaction (**Fig 4.C**). Together, these data reveal that interaction of CHD2 with RTF1 and CHD4 with either GATAD2B or ADNP are likely be mediated by the APRs located in their C-termini.

### APR contributes to CHD4 interaction with NuRD and ChAHP complexes

CHD4 is uniquely found in both the NuRD and ChAHP complexes. This suggests that CHD4 may contain a shared interaction interface with GATAD2A/B and ADNP (core components of both complexes respectively). Thus, we hypothesised that the aliphatic stretch located in CHD4 (1735-1742 aa, YWLLAGII), the sole APR predicted in CHD4 C-terminus, could contribute to CHD4 interaction with both ADNP and GATAD2B (**Fig 5.A**). This motif is evolutionarily conserved in CHD4 orthologues and between CHD subfamily II members (**Fig 5.A**). To examine the role of this putative APR in mediating protein-protein interactions we conducted an AP-MS experiment using FLAG-tagged CHD4-C and a deleted APR mutant (CHD4-C-APR^Del^) as bait. Our data demonstrated a pronounced enrichment of canonical NuRD subunits and ADNP with CHD4-C, compared to CHD4-C-APR^Del^ mutant (**Fig 5.B**). CHD4 expression levels were similar in both conditions, as highlighted in the centre of the volcano plot. This is also evident in the equivalent amount of unique CHD4-C peptides identified in both experiments, whereas there is a noticeable decrease in the number of identified peptides for all NuRD subunits and ADNP (**Fig 5.C**). To further corroborate these observations, we performed pairwise co-immunoprecipitation of haemagglutinin (HA) epitope-tagged GATAD2B-C versus either FLAG-CHD4-C or -CHD4-C-APR^Del^, as GATAD2B is a direct binding partner of CHD4 within the NuRD complex(Sharifi Tabar *et al*., 2022a; Torrado *et al*., 2017). Our data confirmed the expected reduction in GATAD2B-C protein binding to CHD4-C-APR^Del^ as compared to CHD4-C (lane 2 vs lane 1). Complete abrogation of GATAD2B from CHD4 might be achieved by mutating residues within the other potential interaction interface (1453-1513 aa) predicted by AlphaFold Multimer (see **Fig 4.A**). Together, our data demonstrate that the APR region is important for the physical interaction of CHD4 with GATAD2A/B (**Fig 5.C**).

**Figure 5.**
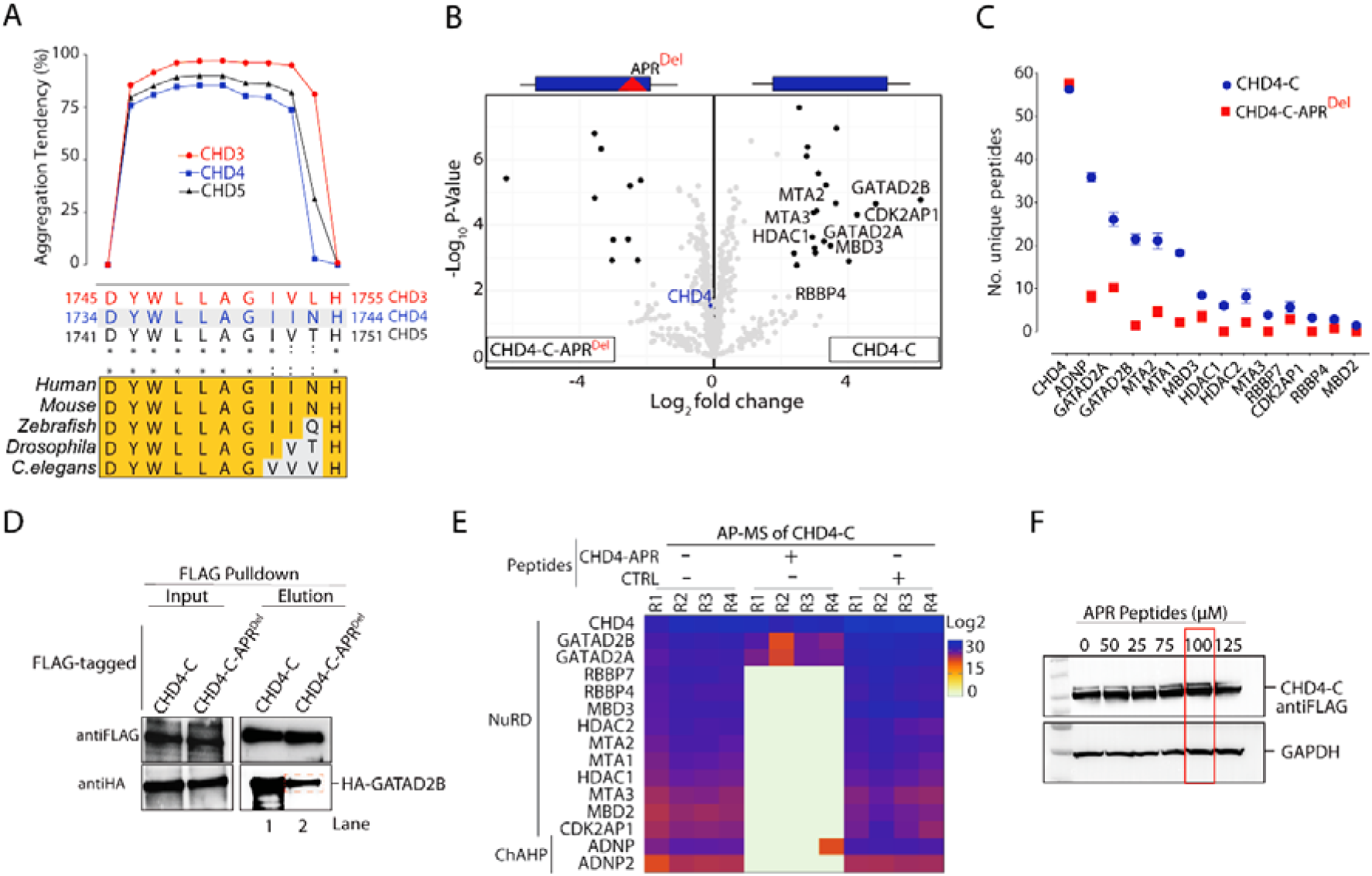
The C-terminus of CHD4 protein harbors an APR region that directly interacts with the NuRD and ChAHP complexes. **A**) TANGO analysis shows β-sheet aggregation tendency for the APR within CHD3, CHD4, and CHD5. Corresponding APR residues are shown with homology within CHD3-5 subfamily and between CHD4 orthologues. **B-C)** AP-MS for interactors of CHD4-C and CHD4-C-APR^Del^ showing: B) volcano plots of enriched proteins; and **C)** The number of unique peptides of subunits of the NuRD and ChAHP complexes detected. **D)** Western blots of input and elution samples from FLAG-CHD4-C co-expressed with HA-GATAD2B protein using an *in vitro* transcription/translation system; samples were probed with anti-HA and anti-FLAG antibodies. **E)** Cellular lysate expressing FLAG-CHD4-incubated in the presence of CHD4-APR or CTRL peptides (100 μM) before subsequent AP-MS. The heatmap represents peptides intensity for of CHD4-NuRD and -ChAHP complex subunits. **F)** Western blots confirming consistent detection of FLAG-CHD4-C protein after addition of APR peptidomimetics (0–125 μM) to the cell lysates. Red box indicates the concentration used in (E).

To eliminate the possibility that the disruption of CHD4-C protein interaction upon deletion of the APR was due to protein misfolding, we adopted an alternative approach using peptidomimetics. Peptides containing the aliphatic stretch (referred to as CHD4-APR) or a stretch of alanine residues (referred to as control (CTRL)), both conjugated to HIV-1 Tat nuclear localisation signal (NLS) were used. In a competition assay, we overexpressed FLAG-tagged CHD4-C in HEK293 cells and mixed the cell lysate with either CHD4-APR or CTRL peptides, followed by AP-MS experiments. Our results demonstrated the complete depletion of the majority of NuRD subunits and ADNP1/2 proteins (**Fig 5.E**). Consistent with the Co-IP results, the enrichment of GATAD2B was diminished but was not completely disrupted. In addition, incubation of FLAG-CHD4-C-expressing cell lysates with a range of CHD4-APR peptide doses, did not diminish FLAG-CHD4-C protein expression (**Fig 5.F**). This observation supports the hypothesis that the CHD4 C-terminal APR contributes to the integration of CHD4 into the NuRD and ChAHP complexes.

### The APR region is critical for the function of the NuRD and ChAHP complexes

As CHD4 is known to regulate gene expression during erythropoiesis, we used mouse G1E-ER4 cells to model cellular differentiation to erythrocytes following induction of GATA-1 expression using tamoxifen(Rylski *et al*, 2003). We first examined the impact of APR peptidomimetics on undifferentiated G1E-ER4 cells using MTT and cell apoptosis assays. Our results showed no significant differences in cell viability or apoptosis (**Supplementary Fig 3.A & B**). To investigate how dissociation of CHD4 from the NuRD and ChAHP complexes affects the gene transcription program during erythropoiesis, we treated G1E-ER4 cells with either CHD4-APR or CTRL peptides (10 μM), followed by induction of GATA-1 expression and performed RNA-Seq analysis (**Fig 6.A**). Successful differentiation of G1E-ER4 cells into erythrocytes following induction of GATA-1 was confirmed by increased expression of erythrocytic signature genes (e.g., *Hba-a1*, *Hba-a2* and *Hba-x*) compared to untreated G1E-ER4 cells (**Fig 6.B**). Differential gene expression analysis of CHD-APR-treated cells over CTRL peptide-treated cells revealed significant dysregulation of 248 genes (**Fig 6.C**). K-means clustering of differentially expressed genes identified two distinct clusters (**Fig 6.C, Supplementary Table 3**). Gene ontology (GO) analysis of cluster 1 featuring downregulated genes upon CHD4-APR treatment revealed enrichment of terms such as glucose catabolic processes and metabolites and energy producing processes. These processes ensure erythrocyte viability and function by providing energy, supporting lipid metabolism, and maintaining erythrocyte integrity. GO analysis of cluster 2 featuring upregulated genes revealed marked enrichment of terms including transcriptional regulation and RNA metabolism (**Fig 6.C**). Taken together, our data reveal a critical interaction hub that facilitates and orchestrates a complex gene expression program leading to normal red blood cells production and metabolism.

**Figure 6.**
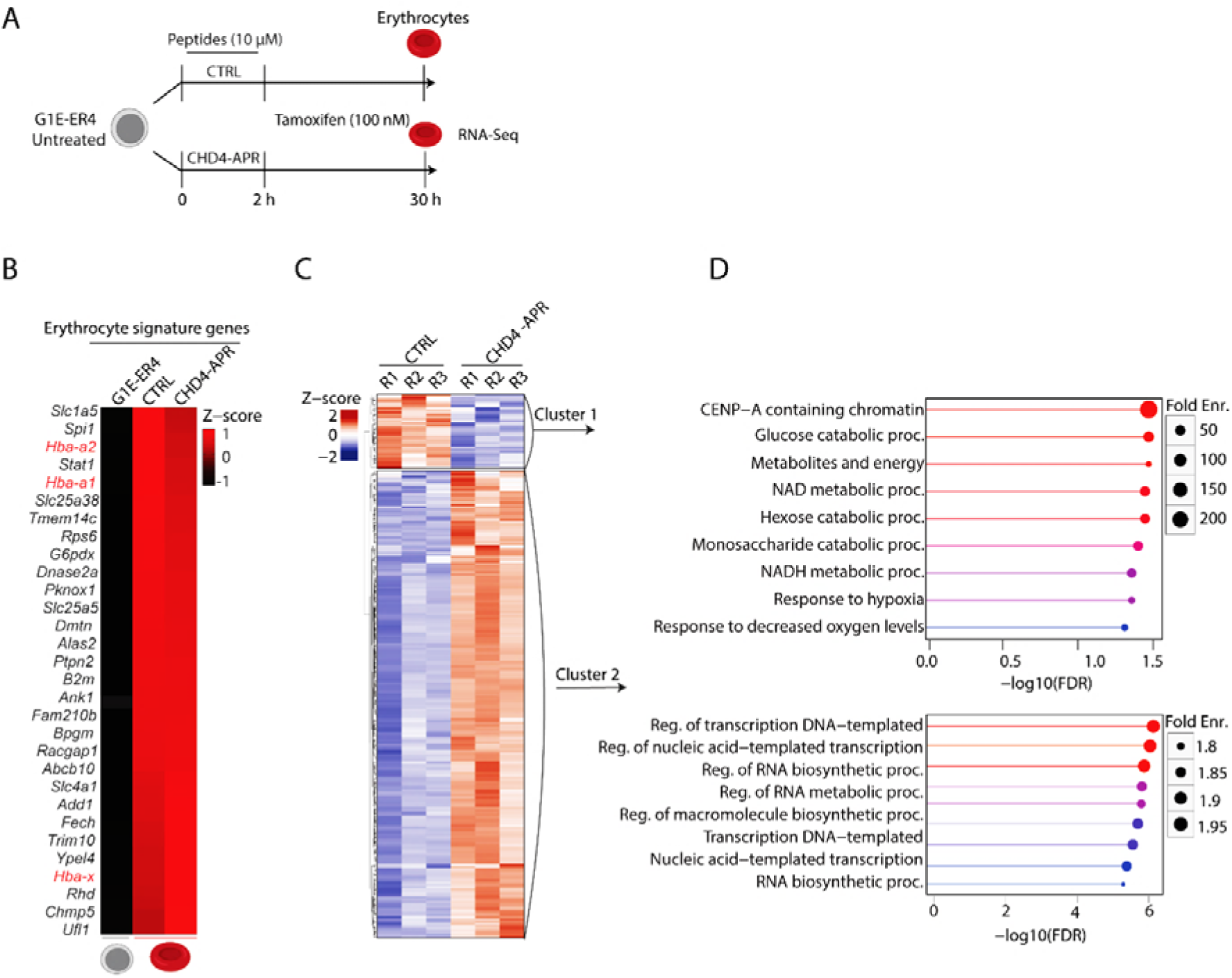
CHD4 dissociation from the NuRD and ChAHP complexes can lead to gene dysregulation. **A)** Schematic illustrating the experimental procedure and sample collection timepoints. G1E-ER4 cells were treated with either CHD4-APR or CTRL peptides for 2 h and then tamoxifen was added, and samples were collected after 28 h (total 30 h). Erythrocytes were subjected to RNA-Seq analysis. **B)** Heatmap representing the upregulation of the erythrocytic signature genes in the presence of tamoxifen (top 30 genes are represented) and globin genes are highlighted in red. RNA-Seq data of untreated G1E-ER4 cells was obtained from ENCODE (GSE101195). The columns represent the mean of combined normalised expression from three replicates. **C)** Z-score heatmap representing the pattern of differentially expressed genes in CHD4-APR vs CTRL peptides. **D)** Gene ontology analysis indicating biological processes significantly associated with the differentially expressed genes.

## Discussion

As CHD proteins are large (ranging in size from 197 to 336 kDa) and contain DNA and histone binding domains (such as ATPase and chromo domains), mapping protein interactions is labour intensive. Past attempts have often resulted in poor data quality. To overcome this issue, we focused on the N- and C-termini of CHD family members and excluded the ATPase and chromo domains to minimise identification of indirect DNA- and histone-mediated protein interactions. The AP-MS approach typically requires the overexpression of bait proteins, but this may lead to their misexpression and/or mislocalisation and thereby increasing the chance of false positives. To address these challenges, we first conducted nuclear fractionation to minimise interaction artifacts arising from protein mislocalisation. Second, we maintained the expression levels of CHD proteins within a similar range by controlling DNA concentration and the number of transfected cells, as confirmed by iBAQ analysis. Additionally, indirect protein interactions bridged by DNA or RNA were minimised by sonicating samples to shear genomic DNA and treating the lysates with nucleases. Detecting known physical interactions, such as those between CHD subfamily I with RTF1(Simic *et al*., 2003), and CHD3/CHD4/CHD5 interaction with the NuRD and ChAHP complexes, verifies our AP-MS workflow and reinforces the potential of our approach to identify novel binding partners.

We utilised Alphafold Multimer in conjunction with our AP-MS data to characterise novel protein complexes. For several of these complexes, we observed high-confidence PAE scores indicating a strong likelihood of direct interactions. However, the low confidence pLDDT scores assigned to intrinsically disordered portions of CHD suggest that Alphafold is limited in its ability to confidently predict the structural details of these interactions, which agrees with previous reports(Bret *et al*, 2024). We also note that the N- and C-termini of CHD proteins may adopt a tertiary structure under physiological conditions, particularly when CHD proteins engage with nucleosomes. Additionally, it is noteworthy that each protein chain within these complexes might also form homodimers or other higher-order assemblies which are difficult to predict with Alphafold.

The novel CHD-specific protein-protein interactions elucidated in this study provide a foundation for a more in-depth exploration of the role of CHD proteins in chromatin biology, encompassing aspects such as gene expression, DNA replication, and repair mechanisms. The interaction of ANP32A, a chaperone crucial for H2A.Z deposition, with CHD6 suggests a role of this complex in the loading of H2A.Z in the regulatory regions of target genes(Mao *et al*, 2014). The interaction of RSBN1 with multiple CHD proteins suggests functional synergy between CHDs, the histone mark H4K20me2 and DNA replication. H4K20me2 is involved in the recruitment of the origin recognition complex for the initiation of DNA replication(Huang *et al*, 2023). The exclusive interaction of uncharacterised KIAA1671 protein with subfamily II CHDs may suggest a distinct biological role for this complex.

Our study provides clear evidence for the role of a unique C-terminal CHD4 APR in co-ordinating the interaction of key functional proteins such as GATAD2B during erythropoiesis. Furthermore, the use of peptidomimetics as a tool to block or interfere with protein interactions mediated via APR-containing interfaces emphasises the utility of this approach. However, several limitations should be considered. Firstly, in our RNA-Seq experiment we used a low concentration of APR which may have limited the inhibition of CHD4 engagement with the NuRD and ChAHP complexes, thereby limiting the degree of gene dysregulation. Timed collection of cells following tamoxifen induction could identify dynamic or temporal changes in gene expression during erythropoiesis. Despite these limitations, our study establishes intriguing paths for further investigations into the molecular mechanisms underlying CHD4 function.

In conclusion, our study has identified the interaction networks of the CHD family of chromatin remodelers in an unbiased manner. Importantly, our identification of a functional interaction hub within an intrinsically disordered region enhances our understanding of the complex regulatory mechanisms that govern the interplay between chromatin remodelers, chromatin dynamics, and gene expression.

## Materials and Methods

### Plasmid constructs

All human orthologues of CHD protein fragments were constructed as GeneBlocks (Integrated DNA Technologies) and were cloned into pcDNA3.1(+) mammalian expression vectors. Each CHD fragment was tagged at the N-terminus with the FLAG epitope. All vectors are available on request.

### Design of APR peptides

CHD4-APR was conjugated to HIV-1 Tat protein nuclear localisation signal (YGRKKRRQRRR) to enhance cell permeability. A peptide representing a stretch of seven alanines was also conjugated to Tat and used as a control. Peptides were synthesised to at least 85% purity by HPLC at Mimotopes, Australia.

### Cell Culture

G1E-ER4 is an erythroid progenitor cell line derived from murine embryonic stem cells that stably expresses a fusion protein combining GATA-1 fused to the estradiol receptor ligand binding domain(Rylski *et al*., 2003). Addition of tamoxifen (100 nM, 4-OHT, Sigma) to the cell culture medium results in the nuclear translocation and transcriptional activation of GATA-1, triggering erythroid differentiation over 1-2 days. G1E-ER4 cells were cultured in Iscove’s Modified Dulbecco’s Medium (500 mL), supplemented with 15% FBS, glutamine, penicillin/streptomycin, CHO Kit ligand-conditioned medium (3 mL, generated in-house from CHO-KLS cells), 1-Thioglycerol (6.2 μL, M6145, Sigma) and mouse erythropoietin (EPO, carrier-free, 2 U/mL, #587608, BioLegend). Cells were maintained in a humidified incubator with 5% CO2 at 37 °C. Expi293F™ cells were cultured to a density of 1.5 × 10^6^ cells/mL in Expi293™ Expression Medium (Thermo Fisher Scientific, USA) on a horizontal orbital shaker (130 rpm).

### Transfection and in vitro transcription/Translation

Polyethylenimine (PEI) was used as a transfection agent (Polysciences, Warrington, PA, USA). The DNA (4 μg) was initially diluted in 200 μL of PBS and then PEI (8 μL, 1 mg/mL) was added, followed by vortexing and then incubation for 20 minutes at room temperature. The mixture was added to the cells cultured in a 12-well plate. Cells were then incubated with shaking for ∼72 h at 37 °C, 5% CO2, in a humidified incubator. For the Co-IP experiment, the TNT® Quick Coupled Transcription/Translation System was used (Promega, #L1170).

### APR treatment for aggregation analysis

Cell pellets were lysed using a buffer (500 μL) consisting of 50 mM Tris-HCl, 150 mM NaCl, 0.5% (v/v) IGEPAL, pH 7.5, 1x protease inhibitor cocktail (Sigma-Aldrich), 1 mM DTT, and 1 μL Pierce™ Universal Nuclease (Thermo Fisher Scientific). Subsequently, the cells were sonicated for 5 cycles, 1 min ON/30 s OFF. Following sonication, the total lysate was collected, and the remaining lysate was portioned into new 1.5 mL Eppendorf tubes. A known concentration of APR-containing peptides was added to each tube, and the mixture was incubated for 30 min at 4 °C. The tubes were then centrifuged at 20,000 g for 30 min to separate the soluble and insoluble fractions.

### Apoptosis assay

G1E-ER4 cells (2.5×10^5^) were treated with 25 µM of CHD4-APR or control APR peptides for 2 h. Cells were then treated with tamoxifen (100 nM) for 28 h followed by labelling with Annexin-APC (BD Biosciences, #550475) and DAPI as per manufacturer’s instructions (Thermo Fisher Scientific). Flow cytometry was then performed to determine the percentage of live, apoptotic or dead cells on an LSR Fortessa (BD Biosciences).

### Sample preparation for nano-LC-MS/MS analysis

FLAG-tagged CHD protein domain pulldown experiments were conducted in triplicate and 9 replicates of FLAG-containing empty vector pulldowns were used to minimise false positive identification. Nuclear fractions were lysed in 800 μL of the same lysis buffer as described above. Lysates were incubated with 20 μL of FLAG antibody-conjugated beads (Sigma-Aldrich) for 2 h. Subsequently, a series of five washes were performed: three washes with a buffer containing 200 mM NaCl, 50 mM Tris-HCl, and 0.5% (v/v) IGEPAL at pH 7.5; and two washes with the same buffer without IGEPAL. Affinity-purified proteins were subjected to on-bead trypsin digestion using 20 μL of digestion buffer comprising 2 M urea freshly dissolved in 50 mM Tris-HCl, 0.5 mM DTT, 100 ng trypsin (Promega, #L1170), with the incubation carried out at 30-35 °C for 2 h. The beads were then collected, and the supernatant was transferred to LoBind tubes. Following this, the beads were resuspended in 20 μL of 2 M urea containing 10 mM iodoacetamide (IAA) in the dark for 20 minutes. The supernatant was transferred back to the previous tube and incubated at 30 °C for 16 h. On the following day, the tryptic peptides were acidified to a final concentration of 2% (v/v) with formic acid and desalted using StageTips (Thermo Fisher Scientific). LC-MS/MS analysis was conducted using an UltiMate™ 3000 RSLCnano System (Thermo Fisher Scientific) coupled to a Thermo Scientific Q-Exactive HF-X hybrid quadrupole-Orbitrap mass spectrometer with a standard nanoelectrospray source. The mass spectrometer was set to a data-dependent acquisition mode (DDA), where each full-scan MS1 operated in a mass scan range of 300 to 1600 m·z^-1^ at a resolution of 60,000. The top 10 most intense precursor ions were selected for fragmentation in the Orbitrap via high-energy collision dissociation activation in the data-dependent acquisition run.

### MS data analysis

To confirm CHD protein expression, an intensity-based absolute quantification (iBAQ) method was used. This algorithm normalises the total intensity of a specific protein by the number of identified unique tryptic peptides. This normalisation accounts for differences in protein length, as longer proteins are expected to yield more tryptic peptides than shorter ones. Analysis of the raw data was conducted using MaxQuant with standard settings. Carbamidomethyl cysteine and methionine oxidation, were chosen as fixed and variable modifications, respectively. Trypsin was selected as the proteolytic enzyme. The output proteingroups.txt table was further processed manually to remove heat shock, plasma membrane, ribosomal, keratin, mitochondrial and cytoplasmic proteins. Proteins represented by at least 2 unique peptides were set aside for analysis in the R studio package as detailed elsewhere(Shah *et al*, 2020). The Perseus algorithm was employed for imputing missing values, with proteins having two missing values being excluded from the analysis.

### Protein Analysis and Structure Prediction

ColabFold is a freely available, open-source implementation of AlphaFold2, designed to accelerate the prediction of protein structures and complexes. ColabFold integrates the rapid homology search functionality of MMseqs2 with the advanced prediction algorithms of AlphaFold2 or RoseTTAFold and is available at https://github.com/sokrypton/ColabFold. ColabFold employs AlphaFold Multimer an extension of AlphaFold2 to predict protein-protein interaction complexes. Two key metrics are generated from this analysis: the PAE (Predicted Aligned Error) and the pLDDT (predicted Local Distance Difference Test) scores to report confidence of prediction(Evans *et al*., 2022; Jumper *et al*, 2021). In the context of protein interaction, the PAE score serves as a metric to evaluate protein-protein interaction likelihood, with low PAE values indicating a high likelihood of protein-protein interactions. The pLDDT score indicates the confidence in the predicted 3D structure, with high pLDDT values reflecting high confidence in the structural prediction(Evans *et al*., 2022). Predictor of natural disorder regions (PONDR) scores were used to evaluate the intrinsically disordered regions. The CHD protein sequences were analysed using the TANGO algorithm to detect APRs based on beta-sheet aggregation tendency (as a percentage)(Fernandez-Escamilla *et al*, 2004). Default physicochemical parameters were selected. APRs with scores below 20% were not considered further.

### RNA extraction and RNA-Sequencing

G1E-ER4 cells were cultured to a density of 2.5 × 10^5^ in 12-well plates. CHD4-APR (- YWLLAGII-Tat) and CTRL (AAAAAAA-Tat) was added to the culture media, each at 10 μM final concentration. After 2 h cells were treated with 100 μM of Tamoxifen for another 28 h and cells were collected for RNA extraction. Cell pellets were lysed in 1 mL of TRIzol™ Reagent (Invitrogen). After a 5-minute incubation at room temperature, 200 μL of RNase-free chloroform (Sigma-Aldrich) was added followed by vigorous mixing. The samples were further incubated for 15 minutes at room temperature and then centrifuged at 14,000 g for 15 minutes at 4°C to separate the phases. The uppermost aqueous layer was carefully transferred to sterile 1.5 mL Eppendorf tubes, and 500 μL of isopropanol (Sigma-Aldrich) and 1 μL of glycogen (Sigma-Aldrich) were added and gently mixed by inversion. The RNA samples were then incubated overnight at -20°C. On the next day, the samples were thawed and centrifuged at 12,000 g for 20 minutes at 4°C. The supernatant was discarded, and the RNA pellet was washed with 1 mL of 75% (v/v) ethanol. After further centrifugation at 12,000 g for 5 minutes at 4°C, the supernatant was removed, and the RNA pellet was allowed to air-dry. Finally, 30 μL of RNase-free, UltraPureTM distilled water (Gibco) was added, and the RNA samples were stored at -80°C.

### Differential Gene Expression analysis

After determining the raw sequence data quality using FASTQC and adaptor trimming using Cutadapt reads were then aligned to the Ensembl mouse genome (GRCm38, release 86) using the STAR aligner (version 2.7.10a) (Dobin *et al*, 2013) (Kechin *et al*, 2017). Gene expression levels were quantified by generating a count matrix using the FeatureCounts command from the Rsubread package(Liao *et al*, 2019), and differential expression analysis was performed using the DESeq2 package in R(Love *et al*, 2014). Heatmaps were generated using R-shiny app FungiExpresZ(Parsania *et al*, 2023). For gene ontology analysis, ShinyGO (version 0.8) was employed on the subset of differentially expressed genes that demonstrated significant changes (FDR < 0.05) and a log2 fold-change exceeding 1.5(Ge *et al*, 2020). The entire list of differentially expressed genes served as the background for this analysis.

## Data Availability

All raw MS data have been deposited to the ProteomeXchange Consortium via the PRIDE partner repository with the dataset identifier PXD055009. Raw RNA-Seq data have been deposited in the NCBI gene expression omnibus (GEO) under accession number GSE275475.

## Acknowledgements

We gratefully acknowledge the technical assistance provided by the Mass Spectrometry Core Facility at The University of Sydney. We also thank Prof. Joel Mackay from The University of Sydney for generously supplying us with G1E-ER4 cells and the GATAD2B C-terminal plasmid (encoding residues 276-593).

## Author contributions

MST and CGB conceived the study, designed MS experiments, analysed data, and wrote the manuscript. CP performed RNA-Seq data analysis. CG, YF, CM, and RN assisted with cloning constructs, prepared samples for MS experiments and performed western blots and MTT assays. ACHW conducted AlphaFold data analysis, BPD performed the flow cytometry analysis. JEJR reviewed the manuscript and provided intellectual input and scientific discussion.

---

**Supplementary Figure 1.**
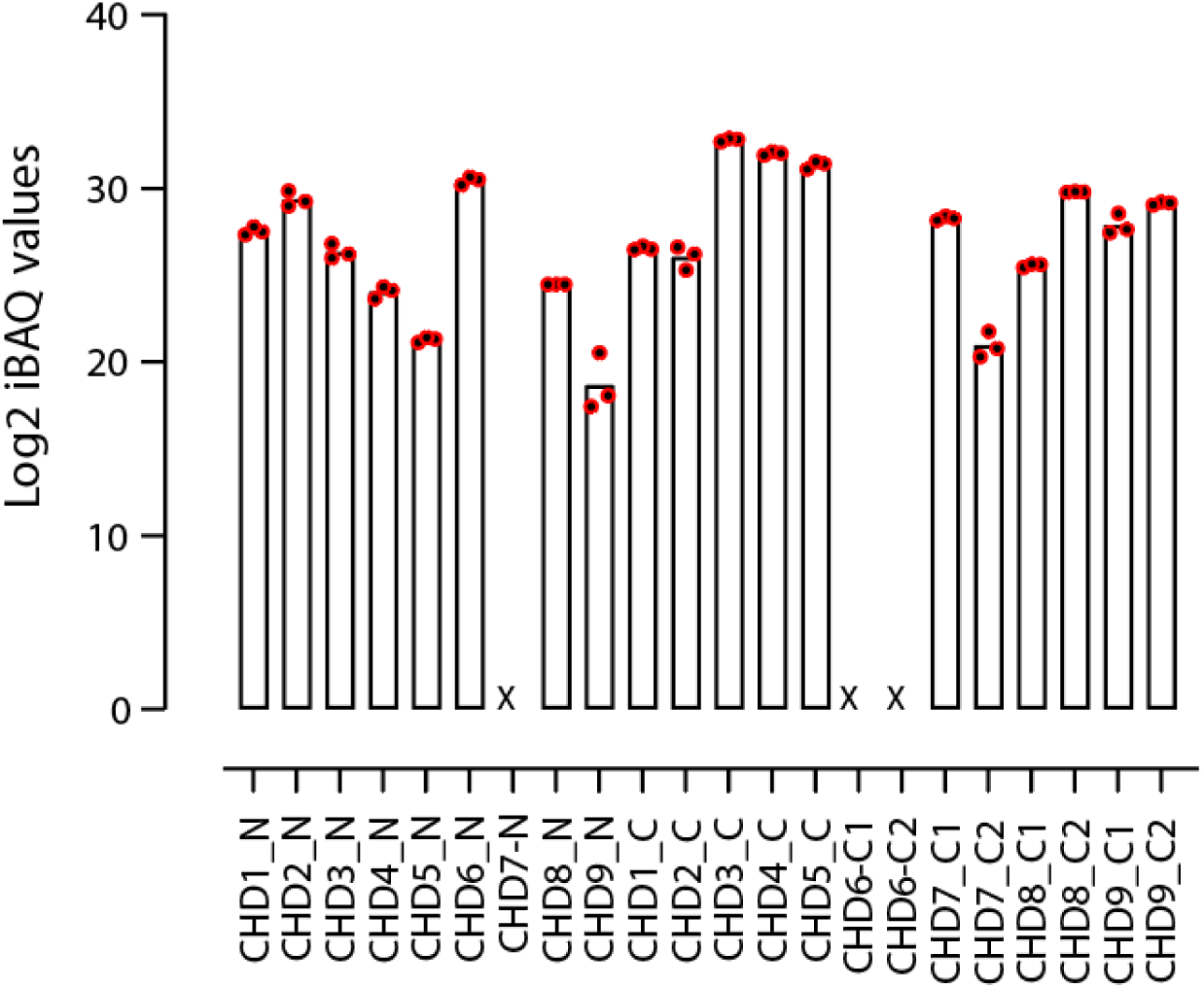
Bar graph showing iBAQ intensities for all FLAG-CHD protein fragments used as a bait in AP-MS experiments.

**Supplementary Figure 2.**
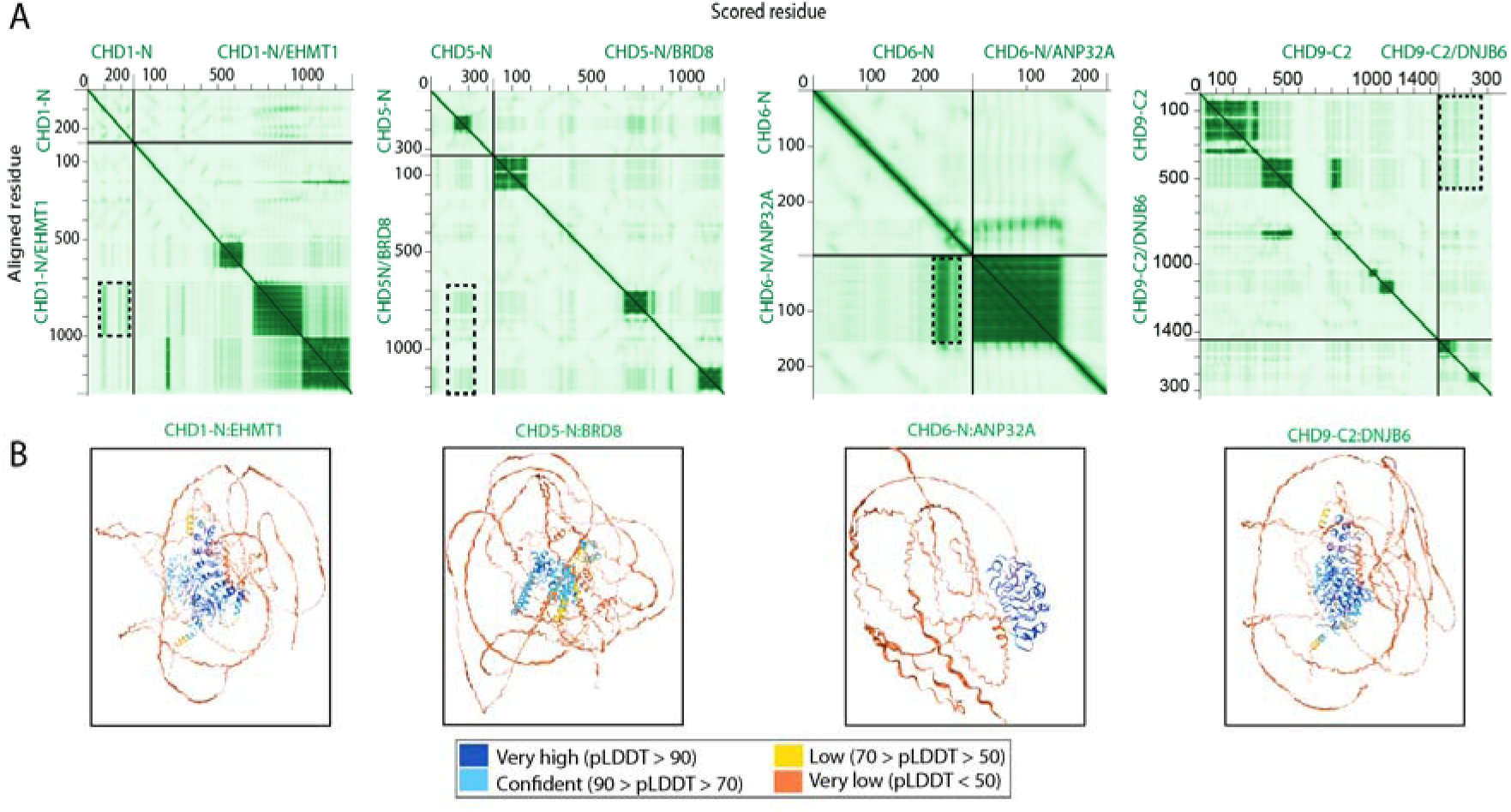
AlphaFold Multimer analysis of potential complexes. **A)** Heatmaps represent the pattern of interaction and potential interaction interfaces highlighted in dotted rectangles. **B)** 3D structures represent the confidence of the predicted structures.

**Supplementary Figure 3.**
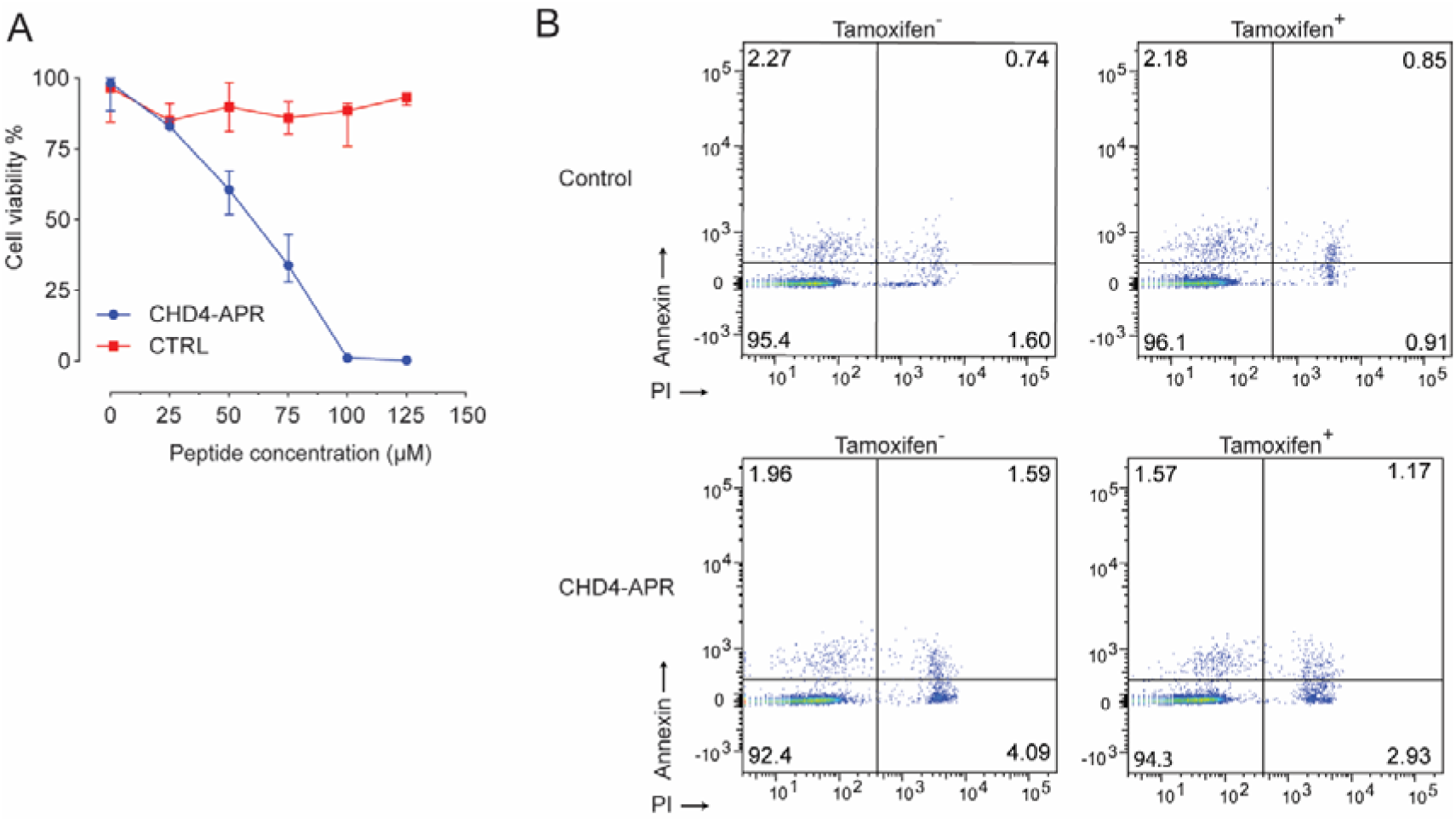
Cell proliferation analysis. **A)** Graph representing the MTT assay performed in three technical replicates (n = 3). Cell viability of G1E-ER4 cells treated with a range of APR peptide concentrations was measured after 48 h. Data are presented as relative to untreated G1E-ER4 cells measured using the MTT assay. **B)** Plot of G1E-ER4 cells treated with 25 μM APR peptides for 2 h, treated with tamoxifen for 28 h, then stained with annexin V and PI and subjected to flow cytometry analysis.

